# Frequency of MicroRNA Response Elements Identifies Pathologically Relevant Signaling Pathways in Cancers

**DOI:** 10.1101/817098

**Authors:** Asha A. Nair, Xiaojia Tang, Kevin J. Thompson, Krishna R. Kalari, Subbaya Subramanian

## Abstract

Complex interactions between mRNAs and microRNAs influence cellular functions. The interactions between mRNA and microRNAs also determine the post-transcriptional availability of free mRNAs and unbound microRNAs. The microRNAs bind to one or more microRNA Response Elements (MREs) predominantly located on the 3’untranslated regions (3’UTR) of mRNAs. In this study, we leveraged MRE sites and their frequencies in transcriptomes of cancer and matched normal tissues to obtain insights into disease-specific interactions between mRNAs and microRNAs. Toward this, we developed a novel bioinformatics method called ‘ReMIx’ that utilizes RNA-Seq data to quantify MRE frequencies at 3’UTR of genes across the transcriptome. We applied ReMIx to The Cancer Genome Atlas (TCGA) Triple Negative (TN) breast cancer tumor-normal adjacent pairs (N=13) and identified distinctly and differentially expressed MREs specific to the TN tumors. Novel data generated by ReMIx identified candidate mRNAs and microRNAs in the MAPK signaling cascade of the TN tumors. We further analyzed the MAPK endogenous RNA network to establish regulatory microRNA partners, along with interacting protein-coding mRNAs that influence and modulate MAPK signaling in TN breast cancers.

## INTRODUCTION

Regulatory interactions between coding and non-coding RNAs in cells determine the post-transcriptional availability of protein-coding mRNA transcripts (Chiang et al. 2010; Guo et al. 2010; Shin et al. 2010; Garcia et al. 2011; Volinia and Croce 2013; Eichhorn et al. 2014; Guo et al. 2014; Lee and Jiang 2017; Rissland et al. 2017; Wu and Bartel 2017). MicroRNAs use seed sequences (6 – 8 bases long) to bind to microRNA Response Elements (MREs) located on the 3’ untranslated regions (3’UTR) of mRNAs. The mRNAs can have one or more distinct MRE sites, thus being targets to multiple microRNAs, and likewise, microRNAs can bind to MRE sites of several different target genes (Krek et al. 2005; Lim et al. 2005). As is known, mRNAs belong to diverse biological pathways, and thus alterations in target gene expression via microRNA binding can impact several cellular processes, such as cell proliferation and apoptosis during cancer development, progression, and migration. Thus, finding critical players among the mRNA-microRNA interacting networks can yield identification of novel therapeutic targets and biomarkers in cancers, especially for cancer subtypes that are least responsive to current modalities of treatment.

Expression profiles of microRNAs and mRNAs (Illumina TruSeq libraries enriched for poly-A RNAs) across many cancer types in The Cancer Genome Atlas (TCGA) were used to infer active and functional microRNA-target interactions in different cancer types (Jacobsen et al. 2013). Alternative polyadenylation of 3’UTRs in TCGA bladder cancer was shown to lead to shortened 3’UTR affecting mRNA stability and attenuated protein translation (Han et al. 2018). Studies have also shown that the presence of SNPs along 3’UTR of genes affect microRNA binding and are associated with multiple cancer subtypes (Pelletier and Weidhaas 2010). However, to our knowledge, no work has been done so far to extrapolate RNA-Seq data to analyze MRE sites and obtain insights into unique interactions between mRNAs and microRNAs at the 3’UTR regions in the tumor and normal-adjacent datasets.

Here we introduce a new bioinformatics approach called ReMIx (pronounced “remix”) – m**R**NA-**M**icroRNA **I**ntegration, that leverages RNA-Seq data to quantify MRE sites at the 3’UTR sequence across the transcriptome. ReMIx profiles MRE sites in tumor and normal samples separately, which later enables identification of differentially expressed MREs that are statistically significant in the tumor samples. Since MRE is the interacting link between mRNAs and microRNAs, ReMIx brings together mRNAs with tumor-specific MRE sites and microRNAs that have the potential to bind to these MRE sites and reports potential mRNA-microRNA candidates that have unique tumor-specific interactions and potential disease-driving functions. This method can be applied to study any cancer type or complex disease with tumor/affected and normal sequencing datasets. To demonstrate the utility of ReMIx, we applied it to the largest RNA sequencing dataset of breast cancer cases and normal-adjacent tissues from TCGA (Cancer Genome Atlas 2012; Ciriello et al. 2015). Using this method, we specifically identified MREs in estrogen receptor positive (ER+), ErbB2 overexpressed–HER2 positive (HER2+), Triple Negative tumors and normal-adjacent tissues. Triple Negative Breast Cancers (TNBC) are highly heterogeneous and one of the most severe forms of breast cancer subtype with no targeted treatment strategies currently available, hence, here we applied ReMIx and identified mRNA-microRNA candidates unique to the TNBC, and not present in ER+ or HER2+ subtypes. Analysis of TNBC data identified MAPK signaling pathway targets as potential biomarkers for TNBC.

## RESULTS

### ReMIx: an automated bioinformatics approach for MRE quantification

We developed an innovative bioinformatics approach called ReMIx, to quantify the expression of MRE sites at the 3’UTR regions of mRNAs using RNA-Seq data. Briefly, ReMIx uses reads aligned to the 3’UTR of genes in a given transcriptome and scans them for evidence of MRE sequences (see Methods for more details). All known MREs for every gene in the reference genome, as reported by TargetScan – human version 7.0 (Agarwal et al. 2015)), are quantified for their level of expression at the 3’UTR of all genes. After quantification, ReMIx normalizes the raw counts of MREs to account for sample library size, 3’UTR length and 3’UTR GC content per gene. Finally, for every gene and for every conserved microRNA that targets the gene, the normalized MRE counts are reported in a tab-delimited format for each gene-microRNA pair in the transcriptome analyzed. The ReMIx workflow is fully automated and designed to run in a multi-threaded cluster environment to analyze paired-end transcriptome samples.

### ReMIx identified 210 Triple Negative breast cancers specific MRE sites

The 3’ untranslated region (UTR) sequences of individual genes (n=12,455, TargetScan v7.0 (Agarwal et al. 2015)) were obtained using the reference human genome hg19 build. Reads aligned to these 3’UTR sequences were obtained using the TCGA Breast Cancer transcriptome dataset for 13 pairs (Tumor and Normal-Adjacent) from TNBC subtype, 56 pairs of ER+ and 20 pairs of HER2+ subtypes, which were then provided as input to the ReMIx workflow (see Methods). The pre-computed MRE sequences (n=329, TargetScan 7.0) were also provided as input to ReMIx to count reads mapped to individual MREs located on each gene. The raw MRE counts were then normalized by factoring library size, 3’UTR lengths and 3’UTR GC content of individual genes. MRE quantification process identified normalized counts for a total of 111,522 MRE sites in tumor and normal adjacent samples separately, for each subtype (Supplementary files 1, 2 and 3 for TN, ER+, and HER2+ respectively).

Next, ReMIx results were used to identify MRE sites that had unique and significant levels of expression (high or low) in TNBC tumors compared to ER+ tumors, HER2+ tumors as well as TN, ER+, and HER2+ normal-adjacent cases. Dunnett-Tukey-Kramer pairwise multiple comparison statistical test was applied to the tumor and normal-adjacent cases across all subtypes (6 groups in total) to highlight MREs that were distinct only to TNBC (p-value < 0.05) when compared to the other two subtypes and all normal-adjacent cases. This resulted in identifying 614 MREs distinct to TNBC (Supplementary file 4). Additionally, the edgeR bioinformatics package (Robinson et al. 2010) was applied to identify differentially expressed MREs by comparing 13 TN tumor and the respective 13 normal-adjacent cases (FDR < 5% and log2FC |2|), and reported 3,053 significant and differentially expressed MREs (Supplementary File 5). By adopting the approach of taking the intersection of MREs reported to be statistically significant and differentially expressed by the two complementary approaches, i.e., DTK (n=614 MREs) and edgeR (n=3,053 MREs), we identified a common set of 210 TN tumor-specific MRE sites. The 210 TNBC MREs are provided in Supplementary file 6. The distinct expression profile of these MRE sites in TNBC with respect to other subtypes and normal-adjacent cases is shown in the heatmap in Figure 1.

**Figure 1.**
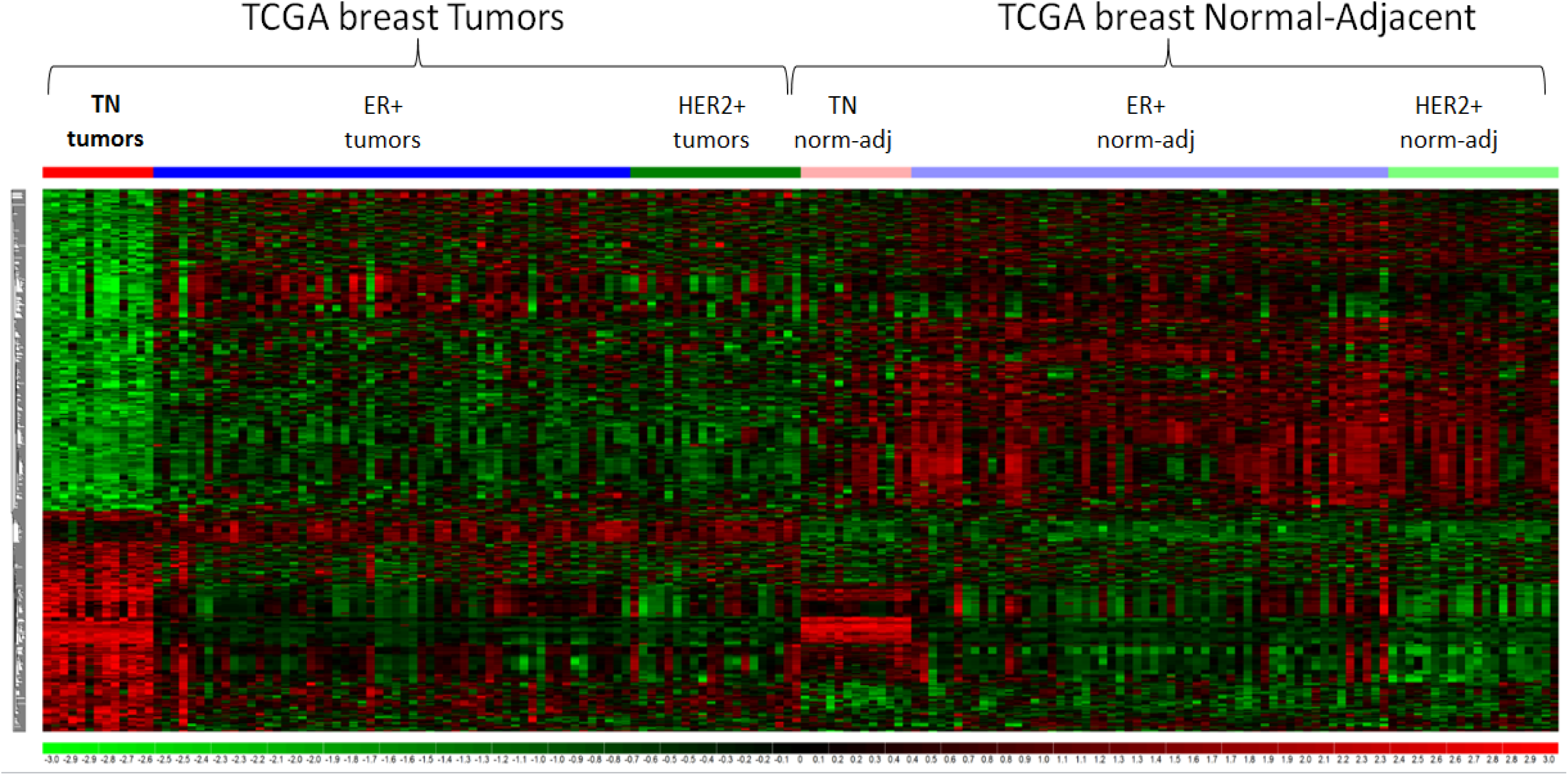
Heatmap of 210 TN-tumor specific MREs. The normalized CQN values of 210 MREs were obtained for TN, ER+ and HER2+ tumors and normal-adjacent (norm-adj) cases. As shown in the heatmap, these MREs have a distinct expression in TN tumors in comparison to the other subtypes as well as TN normal-adjacent cases.

### TNBC relevant MREs are associated with 82 mRNAs and 122 microRNAs

The unique feature of MRE is that it is the interactive site between mRNA and microRNA. Hence, for MRE sites of interest, we can easily decode and obtain information about the mRNA and its interacting microRNA simply by identifying the relevant MREs and decoupling them into their respective mRNA and microRNA pairs. Thus, for the TNBC tumor-specific MRE sites, we deciphered such information for the 210 MREs and obtained a total of 82 mRNAs and 122 microRNAs. From these numbers of mRNAs and microRNAs resulting from the ReMIx analysis, we can construe that over half of the mRNAs (48 out of 82) were used repeatedly and that these mRNAs had multiple MREs on their 3’UTR which were used as interactive sites by different microRNAs.

Next, to further evaluate the mRNAs and microRNAs identified by ReMIx, we analyzed their predominance in terms of differential expression within the respective gene expression and microRNA expression data of the 13 TNBC tumor and normal-adjacent pairs and later, identified the canonical pathways that were associated with the 82 mRNAs and 122 microRNAs.

The differential expression analysis using RNA-Seq data for the 13 TNBC tumor and normal-adjacent pairs indicated that a total of 2,250 genes were differentially expressed (edgeR package (Robinson et al. 2010); statistical significance threshold at FDR < 5% and log2FC |2|). Interestingly, out of the 82 mRNAs identified by the ReMIx analysis, we found that 68 (83%) are also differentially expressed at the gene level between TNBC cases, indicating the high likelihood of microRNA-mediated regulation of gene expression, resulting in genes being significantly differential in expression in the TNBC tumors compared to their normal-adjacent counterparts. Out of the 68 mRNAs, 41 have multiple MREs targeted by different microRNAs. A table listing these 68 mRNAs that are both differentially expressed and have interacting MRE sites for microRNA regulation, as per the ReMIx analysis, can be found in Supplementary file 7.

Likewise, using the microRNA expression data for the 13 TNBC pairs, we found that out of a total of 2,245 microRNAs that were quantified for expression in the tumor and normal-adjacent cases, 778 microRNAs were differentially expressed in the tumor (limma package (Ritchie et al. 2015); adjusted p-value < 0.05). From checking the number of microRNAs identified by ReMIx that was also differentially expressed between TNBC tumor and normal-adjacent, we found that 64 out of 122 microRNAs (52%) were statistically different in expression (FDR < 5%). A table listing the 64 microRNAs that are both differentially expressed and participate in MRE mediated gene expression regulation, as per the ReMIx analysis, can be found in the Supplementary file 8.

Unsupervised hierarchical clustering of the 210 MREs showed that based on the expression profiles of these MREs, the TN cases clustered well within their Tumor and Normal-Adjacent groups. The differential pattern of expression for the 210 MREs is reflected in Figure 2. Notably, unsupervised clustering of the corresponding 82 mRNAs and 122 microRNAs also showed a separation of TNBC into the tumor and normal-adjacent groups, as shown in Figure 2.

**Figure 2.**
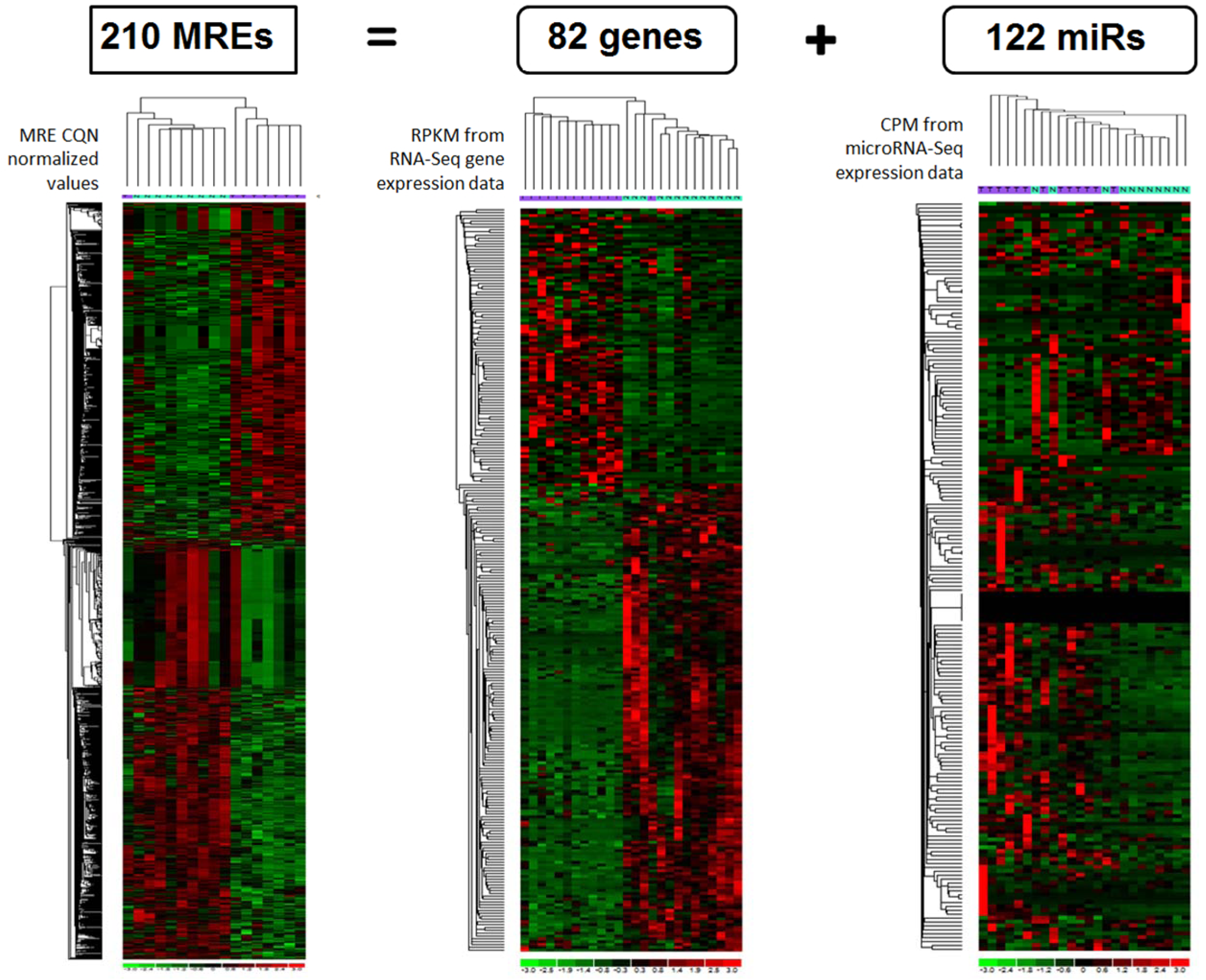
Unsupervised clustering and heatmap representation 210 TN-tumor specific MREs and their associated mRNAs and microRNAs. The 210 MREs were associated with 82 mRNAs and 122 microRNAs. CQN normalized values were obtained for the MRE sites. RPKM normalized values from RNA-Seq and CPM normalized values from microRNA-Seq were obtained for the 13 pairs of TN tumor and normal-adjacent cases. Unsupervised clustering of the cases indicates that tumor (purple) and normal-adjacent (cyan) were clustered well within their groups.

Finally, using RNA-Seq and microRNA differential expression results, the magnitude and direction of change for the 210 MREs and their associated genes and microRNAs in the13 TNBC tumor and normal-adjacent pairs were brought together (Supplementary file 9). We observed that the majority of the MREs follow the same direction as their parent genes, with just a few that do not, likely due to the nature of the TNBC sequencing libraries (Illumina TruSeq).

### MRE associated 122 microRNAs are implicated in TN breast carcinoma

Further analysis of the 122 microRNAs using the TAM 2.0 tool for microRNA set enrichment analysis showed that these microRNAs were associated with cancer pathways as shown in Supplementary Table1. Specifically, 14 out of the 122 microRNAs are also reported in other TNBC studies and are associated with the upregulation of the disease, with an FDR < 2.87e-5. Likewise, 55/122 microRNAs are reported in breast carcinoma studies (FDR < 8.18e-5 and 34/122 in breast neoplasms (FDR < 6.12e-13). Detailed information of these microRNAs is provided in Supplementary Table 1.

### Pathway analysis of 82 genes identified MAPK signaling pathway

The 82 genes obtained from ReMIx were analyzed to identify their associated signaling pathways. Using gene set enrichment analysis (GSEA) (Subramanian et al. 2005) on KEGG and REACTOME databases, the mitogen-activated protein kinase (MAPK) signaling cascade was identified among the top significant pathways. Additionally, the application of the Signaling Pathway Impact Analysis (SPIA) package also confirmed that the MAPK signaling pathway to be activated in TN tumors. The GSEA and SPIA pathway results are provided in Supplementary files 10 and 11 respectively.

Further examination of the genes in the MAPK pathway was conducted by juxtaposing the expression of these genes, obtained from RNA-Seq data of TNBC, with the KEGG-based network of the MAPK pathway. Our analysis revealed that oncogenes KRAS, NRAS, AKT, and NFKB were notably activated and tumor suppressor PTEN was repressed. Figure 3 illustrates the KEGG pathview diagram for the MAPK signaling cascade. MAPK signaling pathway is an extensive cascade with connections to several essential biological pathways downstream such as proliferation, cell cycle, glycolysis, apoptosis, and protein synthesis. As a result, this cascade contains a large number of genes involved in conducting and maintaining activities within the MAPK pathway.

**Figure 3.**
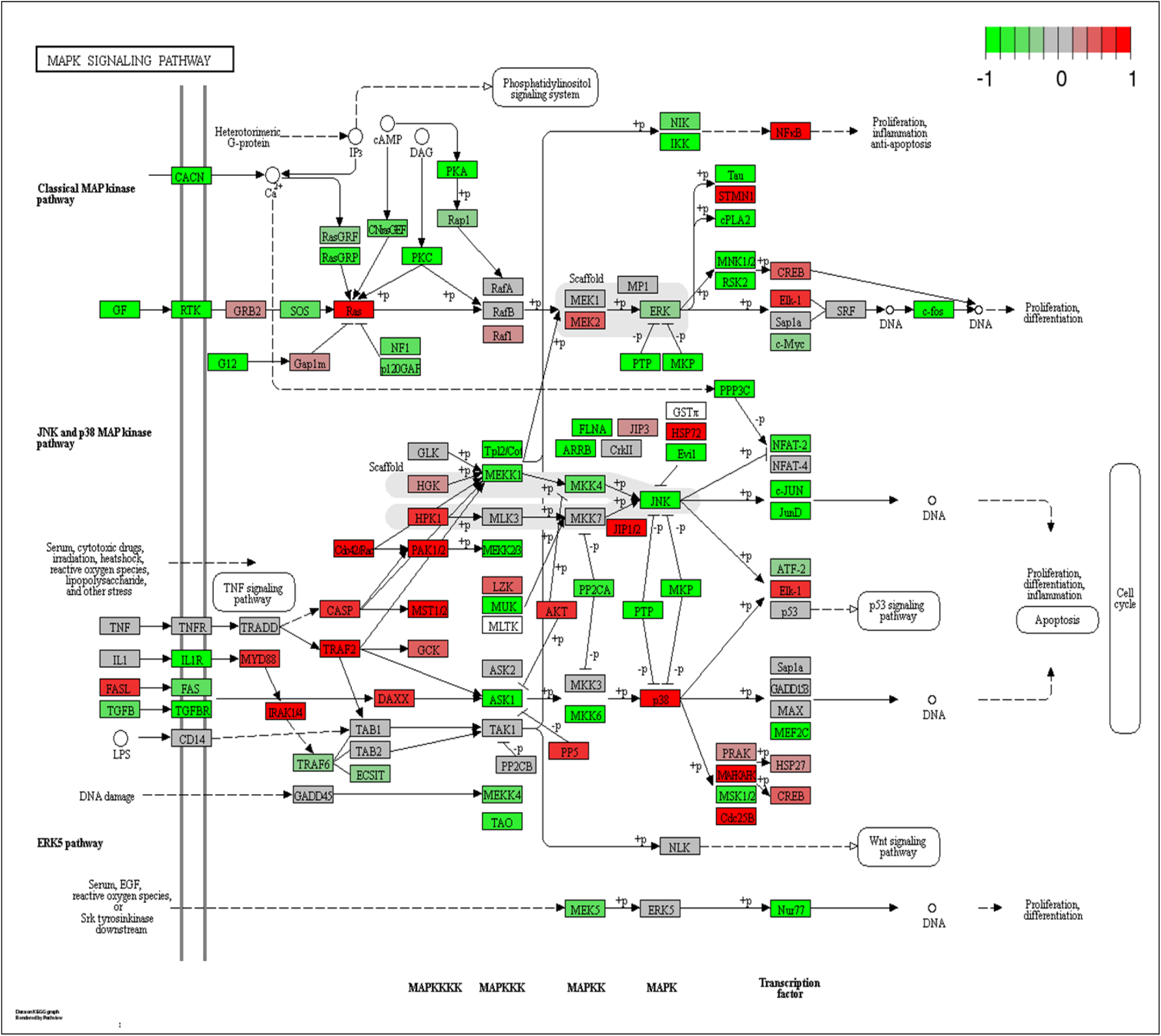
MAPK signaling pathway. Genes in the pathway are colored based on their expression in TN tumors. Oncogenes NFKB and AKT are notably activated in this pathway.

### Expanded MAPK endogenous RNA network including TNBC specific mRNA-microRNA candidates

Based on the MRE results from the ReMIx, we investigated the relevant genes that were associated with MAPK signaling. We found 12 out of 294 genes (~ 4%) in the MAPK pathway have MRE sites with potential for differential binding of microRNAs in TN tumors. The 12 mRNAs with tumor-specific MRE sites and microRNAs with the potential to bind to these sites are provided in Table 1. Next, we expanded the MAPK pathway in the TN tumors by including the interacting microRNAs that, based on our results, are also essential members of these pathways. Figure 4 shows the MAPK endogenous RNA network which represents the genes identified by ReMIx, interacting microRNAs and other protein-coding RNAs that are likely to interact with each other and regulate expressions of the key genes, such as *PI3K, AKT, RAS, NFKB*, and *PTEN*. Taken together, here, we present the expanded network of MAPK signaling cascade and provide a list of potential mRNA-microRNA candidates that interact with each other in this network and could be possible therapeutic targets for TN tumors.

**Figure 4.**
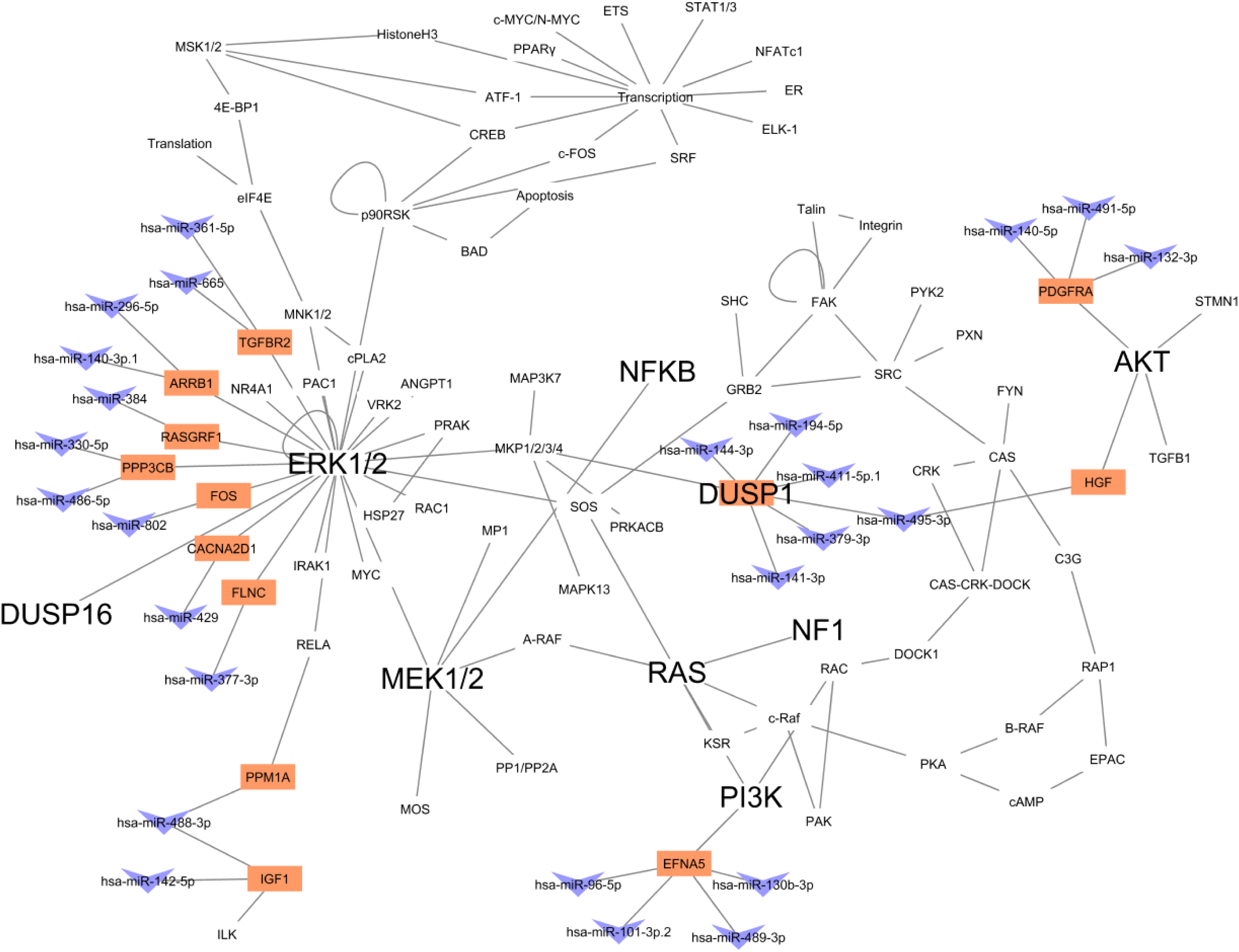
MAPK endogenous RNA network. This figure shows the network of interacting protein-coding mRNAs and non-coding microRNAs in the MAPK singling pathway. The mRNAs and microRNAs reported by ReMIx are represented in colors orange and blue, respectively. Oncogenes AKT, RAS, NF-kB, PI3K, ERK and MEK are shown to interact either directly or indirectly with the mRNA-microRNA candidates.

**Table 1.** Gene-microRNA pairs with distinct TN-specific MRE sites that are part of the MAPK pathway. Table lists genes that are a subset of the 82 genes obtained from ReMIx, and which are members of the MAPK signaling pathway. The microRNAs that bind to the MRE sites which were found to have distinct counts in TN tumors are also provided.

| mRNAs | microRNAs |
| --- | --- |
| **MAPK signaling pathway** |  |
| *CACNA2D1* | hsa-miR-429 |
| *PPP3CB* | hsa-miR-330-5p; hsa-miR-486-5p |
| *RASGRF1* | hsa-miR-384 |
| *IGF1* | hsa-miR-142-5p; hsa-miR-488-3p |
| *HGF* | hsa-miR-495-3p |
| *EFNA5* | hsa-miR-101-3p.2; hsa-miR-130b-3p; hsa-miR-489-3p; hsa-miR-96-5p |
| *PDGFRA* | hsa-miR-132-3p; hsa-miR-140-5p; hsa-miR-491-5p |
| *FOS* | hsa-miR-802 |
| *TGFBR2* | hsa-miR-361-5p; hsa-miR-665 |
| *FLNC* | hsa-miR-377-3p |
| *ARRB1* | hsa-miR-140-3p.1; hsa-miR-296-5p |
| *PPM1A* | hsa-miR-488-3p |

## DISCUSSION

The regulatory interactions between non-coding and protein-coding RNAs have been well established since the past decade, with perhaps the mRNA-microRNA interactions being the most widely studied. While there exists several microRNA target prediction tools such as TargetScan (Agarwal et al. 2015), miRBase (Kozomara et al. 2019), DIANA (Vlachos et al. 2012), PicTar (Krek et al. 2005), miRwayDB (Das et al. 2018), miRanda (Betel et al. 2008), PITA (Kertesz et al. 2007), RNA22 (Loher and Rigoutsos 2012), miRTar (Hsu et al. 2011), not many computational tools have been developed that enable integration of mRNA and microRNA data. MAGIA is a web-based tool for microRNA and genes integrated analysis that brings together target predictions and gene expression profiles using different functional measures for both matched and unmatched samples (Sales et al. 2010). The tool miRmapper uses mRNA-microRNA predictions and a list of differentially expressed mRNAs to identify dominant microRNAs and recognizes similarities between microRNAs based on commonly regulated mRNAs (da Silveira et al. 2018). HisCoM-mimi is a hierarchically structured component analysis method that models biological relationships as structured components, to efficiently yield integrated mRNA-microRNA markers (Kim et al. 2018). These tools use prior knowledge of microRNA target predictions and are developed using unique methodologies to derive mRNA-microRNA interactions. Tools such as miRmapper also have the advantage to highlight the more important microRNAs based on the number of connections it possesses in the studied network. However, the underlying methodologies of all tools are to use the expression of either mRNAs alone or both mRNAs and microRNAs to model their correlation and derive mRNA-microRNA relationships.

With the advent of RNA sequencing (RNA-Seq) technology, profiling of the transcriptome is now possible at base-prevision level. It is a known fact that microRNAs bind to 3’UTR regions of mRNAs to induce their regulatory effects and thereby impact mRNA expression. Studies have shown that shortening of 3’UTR is a frequent phenomenon in cancer to evade oncogenes from microRNA suppression (Xue et al. 2018), repress tumor suppressor genes (Park et al. 2018) and enhance metastatic burden (Andres et al. 2018). Therefore, it is important not only to know which mRNAs are differentially expressed between a tumor and normal pair but also to determine which integration sites or MREs are available along the 3’UTRs of the tumor mRNAs. Identification of MREs that are either present/absent/highly expressed/low expressed in the tumor can potentially shed light on determining the mechanistic nature of tumor progression. While mRNA-microRNA integration tools exist and may be applied to the tumor and normal data, none, to our knowledge, have the ability to precisely report mRNA-microRNA interaction that is solely based on the availability of MREs at the 3’UTR regions. MREs are short 6 – 8 base segments and without appropriate bioinformatics methods, screening RNA-Seq data for MRE sites can yield highly non-specific and erroneous results. This could be a possible reason why this simple, but the highly germane concept has not been explored to date.

In this study, we developed an innovative bioinformatics method called ReMIx that uses RNA-Seq data to identify and quantify microRNA binding sites (commonly known as microRNA Response Elements – MREs) at 3’UTR regions of genes. We applied this methodology to TCGA paired tumor and normal-adjacent breast cancer cases for TN, ER+, and HER2+ subtypes. Using two complementary statistical approaches, we identified 210 MRE sites that have a distinct expression in TN tumor-normal adjacent pairs. Upon de-coupling, we found that the 210 MRE sites corresponded to 82 mRNAs and 122 microRNAs. By reviewing fold changes of these MREs, gene transcripts, and microRNAs, we observed that most of the MREs followed the same direction as their parent genes. We postulate that this is likely driven by the nature of the sequencing library preparation kit (Illumina TruSeq). However, we also found a few MREs with the opposite trend, potentially indicating an alternate 3’UTR mechanism for the gene. Furthermore, we found genes and MREs with positive expression in TNBC tumors but repressed microRNAs, likely indicating the effect of competing endogenous RNAs (ceRNAs) on the microRNAs. Canonical pathway analysis on both mRNAs and microRNAs revealed cancer-related pathways specific to breast cancer. Specifically, the mRNAs identified by ReMIx indicated the MAPK signaling cascade to be significant in TNBC. The mRNAs obtained from ReMIx represented about 4% of gene members in the MAPK pathway. Based on TNBC specific results reported by ReMIx, we expanded the MAPK endogenous RNA network by including the mRNAs with TNBC specific-MRE sites, the corresponding microRNAs that have the potential to bind to these sites, and other protein-coding RNAs in the network that have the potential to interact with each other and regulate expression of primary oncogenes and tumor suppressors in the MAPK signaling pathway. Based on the results from this study, we provide a list of potential mRNA-microRNA candidates that interact with each other at the network level of MAPK signaling cascade and could be possible therapeutic targets for TN tumors.

One of the limitations of this study is that while ReMIx enables identification of candidate mRNA and microRNA players via MRE analysis using RNA-Seq data, this does not establish the fact that the identified microRNAs are indeed present and expressed in the particular disease, in this case, TN tumors. ReMIx results only confirm that the sites on 3’UTR of mRNAs, show distinct expression profiles in tumor and thus have the potential to be regulated by microRNAs in a disease-specific manner. To complement these results, microRNA expression profiles can be used to validate the existence of microRNAs and to check for expression correlation with the corresponding mRNA target(s) identified by ReMIx. This study is limited to microRNA-mediated interactions; however, several other mechanisms modulate gene expression. At the post-transcriptional level, the interplay of other noncoding RNAs, such as long non-coding RNAs, circular RNAs, pseudogenes can collectively form the ceRNA network, and compete with protein-coding genes for microRNA binding, thereby influencing their ultimate impact on gene expression. Also, during transcription, structural and chemical changes such as histone acetylation to determine the accessibility of chromatin domains, and DNA methylation to silence genes, are well-established modes of regulation, especially in cancer.

Lately, there has been much focus in research to explore and identify therapeutic strategies to better treat TNBC patients and improve their chances of survival. It has been shown that activation of the MAPK pathway results in cancer cell proliferation and survival in the tumor (Saini K 2010). Previous studies have shown this pathway to be highly prevalent in TN breast cancer as opposed to other breast cancer subtypes (Hoeflich et al. 2009; Balko et al. 2012; Hashimoto et al. 2014), thus supporting our findings. Studies have also shown that activation of MAPK pathway (Eralp et al. 2008; Gholami et al. 2014; Giltnane and Balko 2014; Hashimoto et al. 2014; Qi et al. 2015; Loi et al. 2016) significantly correlates with tumor proliferation and disease progression in TN tumors. MAPK pathway is a sequentially activated cascade consisting of key genes such as Ras, Raf, MEK, and ERK. Activation of Ras leads to phosphorylation of Raf thereby promoting the activation of MEK and ERK downstream, finally resulting in tumor proliferation and cell survival.

In conclusion, we demonstrate a novel method of using MREs in the identification of functionally relevant mRNA-microRNA interactions that can be potential targets in TNBC. Further, the experimental validation of these interactions is warranted in developing these novel therapeutic targets.

## METHODS

### ReMIx – a novel methodology to compute MRE frequency from RNA-Seq data

We developed an innovative bioinformatics approach called ReMIx, which was used to quantify MRE sites at the 3’UTR regions of mRNAs from RNA-Seq data. ReMIx uses reads aligned to the 3’UTR of genes and scans them for evidence of any given MRE sequence. MRE sequences, which are complementary to the seed sequences of microRNAs, are searched in the 3’UTR of genes that are known to be associated with microRNAs (TargetScan – human version 7.0 (Agarwal et al. 2015)). A hypothetical example of this approach is illustrated in Figure 5. The reads aligned to 3’UTR regions of genes Gene A and Gene B are shown in Tumor and Normal-Adjacent samples. Gene A contains two MRE sites, and Gene B has one MRE site, with a common site (MRE1) in both genes. The number of reads mapped to these MRE sites are quantified for each gene and tabulated separately for Tumor and Normal-Adjacent samples. Later, MRE counts per gene are normalized and statically evaluated to identify differentially expressed MREs for downstream analysis.

**Figure 5.**
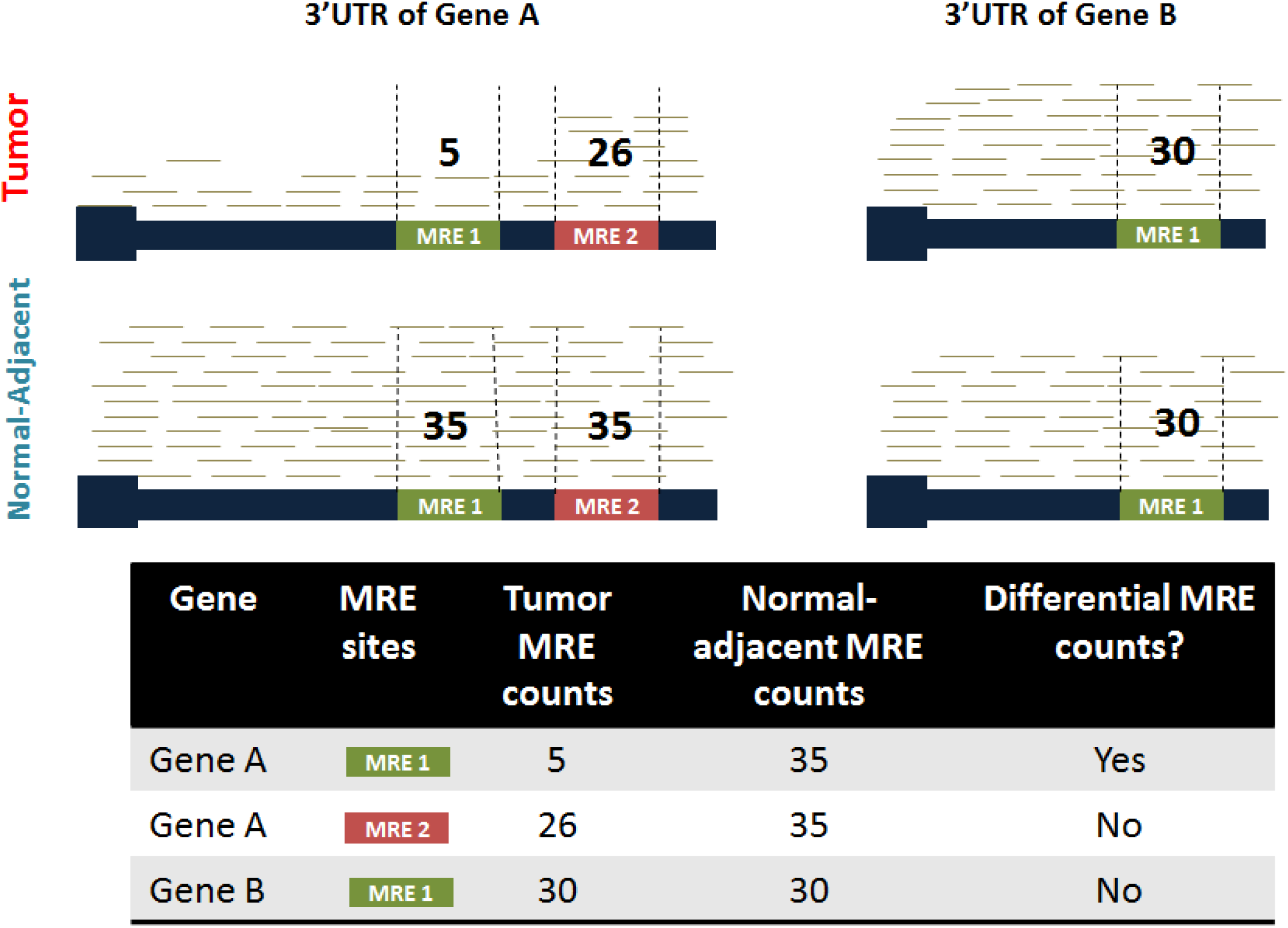
Hypothetical representation of MRE frequency counting using RNA-Seq data. The example shows a Tumor and Normal-Adjacent sample with reads mapped to the 3’UTRs of two genes, Gene A and Gene B that consist of 2 and 1 MRE sites respectively, with a common site (MRE 1). Reads that align with individual MRE sites (vertical dotted lines) are quantified. For every MRE site that belongs to a gene, the counts are then statistically evaluated between Tumor and Normal-Adjacent for evidence of differential frequency (as shown in the table).

### MRE frequency analysis from RNA-Seq data

Seed sequences for all the conserved microRNA families (n=329) were downloaded from TargetScanHuman 7.1, and the corresponding complementary MRE sequences were derived using in-house bioinformatics scripts. Figure 6 is a flowchart representation of the ReMIx methodology. In ReMIx, the RNA-Seq BAM files for both Tumor and Normal-Adjacent were subset to the 3’UTR regions for all genes using the SAMTools suite (Li et al. 2009). The newly obtained BAM files were converted into a FASTQ format for every gene, using the bam2fastx module from Tophat (Trapnell et al. 2009). Next, using MRE sequences of individual microRNAs and the FASTQ files for corresponding genes, the frequency of each MRE was quantified using FIMO (Grant et al. 2011). MRE sites with p-value < 0.05 were selected from the FIMO output for downstream processing. The raw MRE counts were then normalized to account for sample library size, 3’UTR length and 3’UTR GC content per gene. Finally, for every gene and for every conserved microRNA that targets the gene, the normalized MRE counts were reported in a tab-delimited format for each gene-microRNA pair in the Tumor and Normal-Adjacent cases separately.

**Figure 6.**
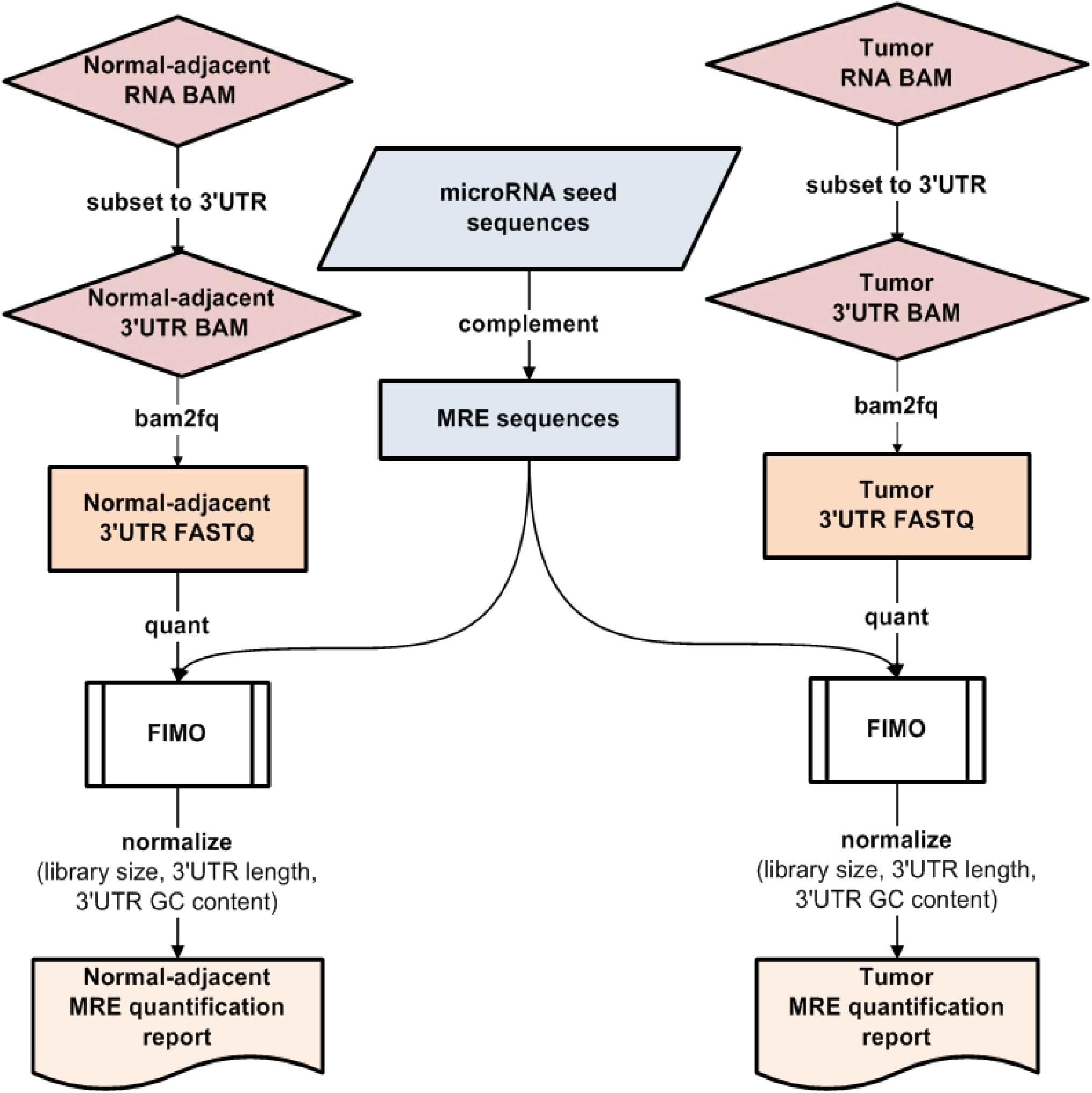
Flowchart representation of MRE frequency quantification from RNA-Seq BAM. The RNA-Seq BAM is subset to 3’UTR regions of all genes, converted to FASTQ and then processed through FIMO to obtain raw MRE counts per microRNA for every target gene. The raw MRE counts are normalized to account for library size, 3’UTR length and 3’UTR GC content and individual Tumor and Normal-adjacent quantification reports are generated.

### 3’UTR definitions obtained from TargetScan

Bartel’s group developed an improved quantitative model to predict canonical targeting of microRNAs to 3’UTR regions of mRNA (Agarwal et al. 2015). A combination of 14 features in the model coupled with experimental approaches such as poly(A)-position profiling by sequencing called 3P-seq was used to define 3’UTR positions of genes in the transcriptome accurately. This data, available at the TargetScan Human 7.1 database, is what was used for 3’UTR definitions of genes in the MRE analysis study.

### RNA-Seq and microRNA-Seq data from TCGA

The RNA-Seq and the microRNA Sequencing fastq files for the TCGA breast cancer samples were downloaded from the TCGA Research Network (http://cancergenome.nih.gov/) using the National Cancer Institute (NCI) Genomic Data Commons (GDC) resource (https://gdc.cancer.gov/). The RNA-Seq fastq files and aligned to the hg19/NCBI 37.1 human reference genome using the MAP-RSeq workflow (Kalari et al. 2014) and the microRNA fastq files were aligned using the CAP-miRSeq workflow (Sun et al. 2014). The normalized microRNA counts from CAP-miRSeq were used to obtain the microRNA expression values in the TNBC samples.

The differential expression analysis of the RNA-Seq data for the TNBC tumor and normal-adjacent pairs were obtained using the bioinformatics R package edgeR (Robinson et al. 2010). The statistical significance threshold used was FDR < 5% and log2FC |2|. For these 13 pairs of TNBC cases, differential expression analysis of the microRNA sequencing data was performed using the R bioinformatics package called limma (Ritchie et al. 2015). The statistical threshold used to identify significantly differential expressed microRNAs was adjusted p-value < 0.05.

### Statistical analyses on MRE sites and activated pathway identification

Evaluation of MRE sites that represented distinct and TNBC-specific expression as opposed to ER+ and HER2+ subtypes and normal-adjacent cases were obtained using the R package Dunnett-Tukey-Kramer Pairwise Multiple Comparison Test Adjusted for Unequal Variances and Unequal Sample Sizes. Statistically significant MRE sites were selected using p-value cut-off < 0.05. The bioinformatics R package edgeR (Robinson et al. 2010) was used to obtained differentially expressed MREs between TN tumors and matched normal-adjacent pairs at FDR <5% and log2FC |2|.

### Pathway analysis for canonical pathways

The microRNA set analysis tool called TAM2.0 was used to identify cancer-related pathways for the 122 microRNAs. Likewise, enriched canonical pathway analysis for 82 genes was performed using KEGG and Reactome functional databases. Open source analysis toolkit WebGestalt (Wang et al. 2017) was also used for pathway identification by using the option to perform Gene set enrichment analysis (GSEA). Identification of the relevance and activation/inhibition status of pathways was evaluated using the R package called Signaling Pathway Impact Analysis (SPIA). The Bioconductor R package called Pathview (Luo et al. 2017) was used to map the gene expression data from TNBC and visualize the MAPK pathway using the KEGG-based network model of this pathway.

## Supporting information

Supplementary summary

## AUTHOR CONTRIBUTIONS

SS and KK conceived the study. AN performed all analyses and wrote the manuscript. KK and SS supervised and helped AN interpret the data. XT and KT helped with subtyping the TCGA breast cancer RNA-Seq data and provided useful suggestions regarding this study. All authors read and approved the final manuscript.

## CONFLICTS OF INTEREST

The authors declare that they have no competing interests.

## FUNDING

This work is supported by the Center for Individualized Medicine (CIM) at the Mayo Clinic in Rochester, MN and funds from the University of Minnesota, Medical School Innovation award and the Department of Surgery, Research funds.

---

**Supplementary Table 1.**
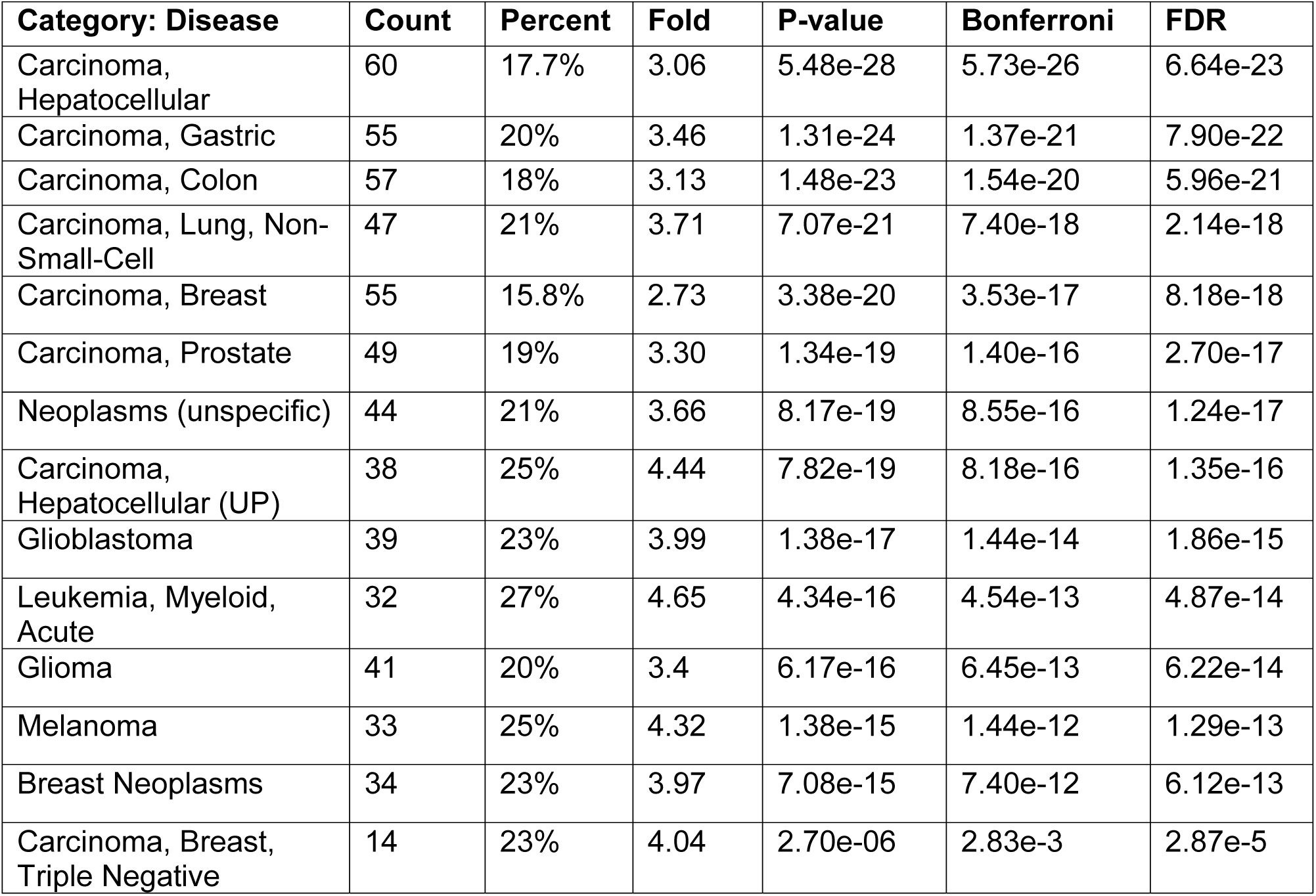
TAM 2.0 results for microRNA set pathway analysis.

| Category: Disease | Count | Percent | Fold | P-value | Bonferroni | FDR |
| --- | --- | --- | --- | --- | --- | --- |
| Carcinoma, Hepatocellular | 60 | 17.7% | 3.06 | 5.48e-28 | 5.73e-26 | 6.64e-23 |
| Carcinoma, Gastric | 55 | 20% | 3.46 | 1.31e-24 | 1.37e-21 | 7.90e-22 |
| Carcinoma, Colon | 57 | 18% | 3.13 | 1.48e-23 | 1.54e-20 | 5.96e-21 |
| Carcinoma, Lung, Non-Small-Cell | 47 | 21% | 3.71 | 7.07e-21 | 7.40e-18 | 2.14e-18 |
| Carcinoma, Breast | 55 | 15.8% | 2.73 | 3.38e-20 | 3.53e-17 | 8.18e-18 |
| Carcinoma, Prostate | 49 | 19% | 3.30 | 1.34e-19 | 1.40e-16 | 2.70e-17 |
| Neoplasms (unspecific) | 44 | 21% | 3.66 | 8.17e-19 | 8.55e-16 | 1.24e-17 |
| Carcinoma, Hepatocellular (UP) | 38 | 25% | 4.44 | 7.82e-19 | 8.18e-16 | 1.35e-16 |
| Glioblastoma | 39 | 23% | 3.99 | 1.38e-17 | 1.44e-14 | 1.86e-15 |
| Leukemia, Myeloid, Acute | 32 | 27% | 4.65 | 4.34e-16 | 4.54e-13 | 4.87e-14 |
| Glioma | 41 | 20% | 3.4 | 6.17e-16 | 6.45e-13 | 6.22e-14 |
| Melanoma | 33 | 25% | 4.32 | 1.38e-15 | 1.44e-12 | 1.29e-13 |
| Breast Neoplasms | 34 | 23% | 3.97 | 7.08e-15 | 7.40e-12 | 6.12e-13 |
| Carcinoma, Breast, Triple Negative | 14 | 23% | 4.04 | 2.70e-06 | 2.83e-3 | 2.87e-5 |

