## Supplementary summary for "Frequency of MicroRNA Response Elements Identifies Pathologically Relevant Signaling Pathways in Cancers"

**Supplementary file 1:** CQN (conditional quantile normalization) values for 111,522 MREs for TCGA TN (Triple-Negative) tumor and matched normal-adjacent samples

**Supplementary file 2:** CQN (conditional quantile normalization) values for 111,522 MREs for TCGA ER+ (Estrogen Receptor +) tumor and matched normal-adjacent samples

**Supplementary file 3:** CQN (conditional quantile normalization) of TCGA ErbB2 overexpressed–HER2 positive (HER2+) tumor and matched normal-adjacent samples

**Supplementary file 4:** 614 MREs with distinct expression in TCGA Triple-Negative tumors identified using Dunnett-Tukey-Kramer pairwise multiple comparison

**Supplementary file 5:** 3,053 significant and differentially expressed MREs distinct to TCGA Triple-Negative tumors using edgeR bioinformatics package

**Supplementary file 6:** 210 MREs that are significant and differential expressed only in TCGA Triple-Negative tumors (common candidates from DTK and edgeR

**Supplementary file 7:** 68 mRNAs that are both differentially expressed and have interacting MRE sites for miRNA regulation

**Supplementary file 8:** 64 miRNAs that are both differentially expressed and participate in MRE mediated gene expression regulation according to ReMIx analysis

**Supplementary file 9:** Direction and magnitude of fold change for 210 MREs and their associated genes and microRNAs

**Supplementary file 10:** GSEA (gene set enrichment analysis) ranked by 68 mRNAs identifed to ReMIx to have interacting MRE sites for miRNA regulation

**Supplementary file 11:** SPIA (signaling pathway impact analysis) on 68 mRNAs identified by ReMIx to have interacting MRE sites for miRNA regulation
